# Comprehensive Genomic Features indicative for Notch Responsiveness

**DOI:** 10.1101/2023.12.04.569922

**Authors:** Benedetto Daniele Giaimo, Tobias Friedrich, Francesca Ferrante, Marek Bartkuhn, Tilman Borggrefe

## Abstract

Transcriptional specificity is often determined by transcription factor levels and/or chromatin context. In the Notch signal transduction pathway, transcription factor RBPJ is the central component and directly forms a coactivator complex together with the Notch intracellular domain (NICD). While the RBPJ protein levels remain constant in most tissues, dynamic expression of Notch target genes varies depending on the given cell-type and the Notch activity state. To elucidate dynamic RBPJ binding genome-wide, we investigated RBPJ occupancy by ChIP-Seq making use of Notch-dependent T cells. Surprisingly, only a small set of the total RBPJ sites show a dynamic binding behavior in response to Notch signaling. Compared to static RBPJ sites, dynamic sites differ in regard to their chromatin state, binding strength and enhancer positioning. Dynamic RBPJ sites are predominantly located distal to transcriptional start sites (TSS), while most static sites are found in promoter-proximal regions. Importantly, gene responsiveness is preferentially associated with dynamic RBPJ binding sites and this static and dynamic binding behavior is repeatedly observed in different cell types and species. Based on the above findings we used a machine-learning algorithm to predict Notch responsiveness with high confidence in different cellular contexts. This approach is potentially applicable to other transcription factors regulating signal-induced gene sets. Our results strongly support the notion that the combination of binding strength and enhancer positioning are indicative of Notch responsiveness.

## INTRODUCTION

Signal transduction pathways enable transmission of extracellular signals, environmental changes or mechano-transduction into changes in gene expression. For several signaling pathways, it is known that ligand or stimulus-dependent changes lead to the translocation into the nucleus of a pathway-specific transcription factor (TF; i.e. SMADs, NFκB). Alternatively, TF levels in the nucleus may remain constant but changes in the associated cofactors occur (i.e. β-catenin/TCF and RBPJ). While changes in gene expression have been already studied using transcriptomics approaches, studies combining TF occupancy combined with dynamic gene expression remain scarce.

In regard to the Notch signaling pathway, ligand binding leads to the release of the Notch intracellular domain (NICD) resulting in the assembly of a coactivator complex, also known as Notch transcriptional complex (NTC) (1,2). This complex consists of TF RBPJ, also known as CSL (*Homo sapiens* CBF1, *Drosophila melanogaster* Suppressor of Hairless, and *Caenorhabditis elegans* Lag-1), together with NICD and, among others, the acetyltransferase EP300 to promote the expression of Notch target genes. In absence of Notch signaling, RBPJ and HDAC-containing associated corepressors inhibit Notch target genes expression. Several studies have focused on the identification and characterization of components of both the corepressor and coactivator complexes, thereby elucidating on the regulation of the chromatin environment at Notch target genes (3–13). The exact repressing mechanism is well-documented and includes additional cofactors such as the SHARP/NCoR complex recruiting HDACs (8,9,12) and histone demethylases (6,10,14). Several studies have also analyzed the genome-wide distribution of RBPJ in various different models (15–25). Although it was previously thought that RBPJ binds to its cognate enhancers regardless of Notch status, more recent studies have revealed a different scenario. In fact, Notch activation significantly increases the binding of RBPJ (15,21,24,26). So far, the functional meaning and the underlying molecular mechanism(s) determining these changes in RBPJ occupancy remained unclear.

Here we determined either “static” or “dynamic” RBPJ sites in a mouse progenitor T cell line with characteristic constitutive Notch signal activity. Only dynamic occupancy of RBPJ correlates with Notch-dependent changes in the chromatin state and gene expression. Dynamic RBPJ occupancy characterizes Notch responsiveness also in human T-ALL cell lines as well as breast cancer cell lines, which allowed us to develop a testable machine-learning approach. Thus, we can identify Notch responsiveness using a machine-learning algorithm solely based on genome-wide occupancy of RBPJ in a variety of cell types.

## MATERIALS & METHODS

### Cell culture and treatments

Mouse leukemia preT cells (Beko) were grown at 37°C under 5% CO_2_ in Iscove’s Modified Dulbecco Medium (IMDM, Gibco 21980-065) supplemented with 2% fetal bovine serum (Pan Biotech), 5 mg/l insulin (Sigma-Aldrich), 0.3 mg/ml Primatone, nonessential amino acids (Gibco) and penicillin/streptomycin (Gibco).

*Drosophila melanogaster* Schneider cells were grown in Schneider’s *Drosophila* medium (Gibco 21720024) supplemented with 10% fetal bovine serum (Gibco 10270-106), Glutamine (Gibco 25030-024) and penicillin/streptomycin (Gibco).

Beko cells were treated with 10 μg/ml GSI (DAPT; Alexis ALX-270-416-M025) or with DMSO as control for 24 hours. In the case of the washout experiments, Beko cells were incubated with 10 μg/ml GSI (supplemented freshly at 24 hours after washing away the old one) for 48 hours. Subsequently, cells were collected, washed and placed in culture for additional 24 hours before performing the experiment.

### Protein extracts and Western blotting

Nuclear extracts were prepared by washing the cells twice in PBS and resuspending them in Hypotonic buffer (20 mM Hepes pH 7.9, 20 mM NaCl, 5 mM MgCl_2_, 10% glycerol, 0.2 mM PMSF). After 20 min incubation on ice, samples were centrifuged (4.000 rpm, 10 min, 4°C) and the nuclei were washed twice with ice-cold PBS. Nuclei were lysed in Hypertonic Buffer (20 mM Hepes pH 7.9, 1 mM MgCl_2_, 300 mM NaCl, 0.2% NP-40, 25% glycerol, 0.2 mM PMSF, 1x Protease inhibitor mix, 0.3 mM DTT) and incubated 20 min on ice. After centrifugation (14.000 rpm, 5 min, 4°C), the supernatants were collected and protein concentration was measured by Bradford assay (Sigma-Aldrich). Extracts were boiled in presence of SDS loading buffer and analyzed by Western blotting.

For Western blotting purposes, proteins were resolved in SDS polyacrylamide gels and transferred to a Nitrocellulose membrane (Amersham) by wet blotting.

Membranes were blocked in 5% milk / TBST (1x TBS, 0.1% Tween 20) and incubated over night with the desired antibody diluted in 5% milk / TBST [1:1000 H3 (abcam ab1791); 1:1000 Val1744 cleaved NICD1 (Cell Signaling Technology 4147)]. Membranes were washed in TBST and incubated 1 hour at room temperature with secondary antibody diluted 1:5000 in 5% milk / TBST [anti-rabbit IgG HRP (Cell Signaling 7074S)]. Membranes were washed in TBST and incubated at room temperature with ECL solution. Chemiluminescence was detected with a Vilber Fusion FX7 system.

### RNA extraction, libraries and sequencing

Total RNA was purified using the RNeasy Mini Kit (Qiagen #74104), the QIAshredder (Qiagen #79654) and the DNase I (Qiagen #79254) accordingly to manufactureŕs instructions. Libraries were prepared using the TruSeq® Stranded Total RNA LT - Ribo-Zero Gold kit (Illumina RS-122-2301/2) and sequenced on a NextSeq device.

### RNA-Seq and microarray analysis

Basic summary statistics for the RNA-Seq experiments are provided in Table S7.

The previously published microarray and RNA-Seq datasets used in this study are indicated in Table S7. Microarray data was downloaded within R v. 4.0.3 with the *getGEO* function of the GEOquery (Davis and Meltzer 2007) package. Since the data were already log normalized, the log_2_FC was calculated by subtracting the treatment values from the control value RNA-Seq analysis was performed within R using a custom-made version of the systemPipeR (27) R/BioConductor package. Raw sequencing reads were adaptor and quality trimmed using trimGalore v. 0.6.5 (https://github.com/FelixKrueger/TrimGalore. The quality of the trimming was validated by visual inspection after using systemPipeR’s *seeFastq* function. Trimmed FASTQ files were aligned against the mouse (mm9) or human (hg19) genome using HISAT v. 2.2.1 (28) with the parameters “–phred33 –k 1 –min-intronlen 30 –max-intronlen 3000” and stored as sequence alignment maps. Conversion from sequence alignment map format to binary alignment map format (BAM) was done using the *runCommandline* function of the systemPipeR package. These BAM files were quality checked by using systemPipeR’s *alignStats* function and subsequently a gene reads table was calculated by the *summarzieOverlaps* function of the GenomicAlignments (29) R/BioConductor package and the corresponding Gene Transfer Format (GTF) file (Illumina’s iGenomes). Normalization and detection of deregulated genes was performed using the DESeq2 v. 1.28.1 package (30). Criteria for the definition of significantly deregulated genes were log2FC > 1 or < −1 and adjusted p-value < 0.05.

### Chromatin immunoprecipitation (ChIP), libraries and sequencing

ChIP experiments were performed as previously described (25). Briefly, cells were fixed in 1% FMA for 30 min at room temperature. The FMA reaction was blocked for 5 min at room temperature by adding 1/8 volume of 1 M glycine pH 7.5. Cells were washed twice with PBS and resuspended in 1 ml of SDS Lysis Buffer (1% SDS, 10 mM EDTA, 50 mM Tris-HCl pH 8.1). After incubation on ice for 10 min, samples were sonicated using a Covaris System S220 AFA (28 cycles, 30 sec ON, 30 sec OFF). The chromatin was diluted in ChIP Dilution Buffer (0.01% SDS, 1.1% Triton X-100, 1.2 mM EDTA, 16.7 mM Tris-HCl pH 8.1, 167 mM NaCl) and pre-cleared with protein-A-Sepharose beads (GE Healthcare 17-5280-02) for 30 min at 4°C. The chromatin was subsequently incubated over night with the proper amount of the desired antibody and the antibodies were immobilized with 40 μl protein-A-Sepharose beads for 1 hour at 4°C with rotation. Depending on the antibody, different combinations of the following washing buffers were used: low salt washing buffer (0.1% SDS, 1% Triton X-100, 2 mM EDTA, 20 mM Tris-HCl pH 8.1, 150 mM NaCl), high salt washing buffer (0.1% SDS, 1% Triton X-100, 2 mM EDTA, 20 mM Tris-HCl pH 8.1, 500 mM NaCl), LiCl washing buffer (1% NP-40, 1 mM EDTA, 10 mM Tris-HCl pH 8.1, 0.25 M LiCl) and TE buffer (10 mM Tris-HCl pH 8.0, 1 mM EDTA). Chromatin was eluted from beads with Elution Buffer (50 mM Tris-HCl pH 8.0, 10 mM EDTA pH 8.0, 1% SDS) and crosslinks were reverted at 65°C over night. After diluting the SDS with TE buffer, samples were incubated with RNase A (Thermo Scientific EN0531) at 37°C for 2 hours and subsequently with Proteinase K (Invitrogen 25530-049) at 55°C for 2 hours. After extraction with phenol*/*chloroform/isoamylic alcohol, the DNA was purified using the Qiaquick PCR cleanup kit (Qiagen 28104).

The following antibodies were used: H3K4me1 (abcam ab8895), H3K4me3 (Diagenode pAb-003-050), or RBPJ (Cell Signaling Technology 5313).

ChIP experiments were analyzed on a StepOnePlus^TM^ sequence detector system (Applied Biosystem) using specific oligonucleotides and double-dye probes (Table S9).

In the case of the ChIP-Seq, chromatin from *Drosophila melanogaster* Schneider cells was used for spike-in purposes (each 25 μg of mouse chromatin, 50 ng or 25 ng of *Drosophila* chromatin were used in ChIP versus histone proteins or TFs respectively). 2 μg of anti-His2Av (Active Motif 61686) were added to each immunoprecipitation for spike-in purposes. Libraries were prepared using the Diagenode MicroPlex Library Preparation kit v2 (Diagenode C05010012) or the Diagenode MicroPlex Library Preparation kit v3 (Diagenode C05010001) following manufacturer’s instructions with few modifications. Libraries were purified with Agencourt AMPure XP Beads (Beckman Coulter, #A63881), quantified, analyzed on a Tapestaion device (Agilent) and pooled. Finally, sequencing was performed on a HS2500 or a NovaSeq device.

### ChIP-Seq analysis

Basic summary statistics for the ChIP-Seq experiments are provided in Table S7.

The previously published ChIP-Seq datasets used in this study are indicated in Table S7. Primary processing of raw sequencing reads until generation of BAM files was done as described for the RNA-Seq analysis. HISAT2 was used for both single- and paired end reads with ChIP-Seq specific parameters “-k 1 –no-spliced-alignment –phred33”. Duplicated reads were removed from BAM files using Picard Tools (available at http://broadinstitute.github.io/picard/) MarkDuplicates function with additional parameters “REMOVE_SEQUENCING_DUPLICATS=true REMOVE_DUPLICATES=true”. MACS2 v. 2.2.7.1 (31) was used with a q-value threshold of 0.01 to call peaks for the individual BAM files with or without input. When ChIP-Seq replicates were available, MSPC v.4.0.0 (32) was used with parameters “-r bio -w 1e-6 -s 1e-10” to validate the called peaks and determine the set of “true positive peaks”. Only if a true positive RBPJ peak was conserved in 3 out of 5 control replicates (Beko) or 2 out of 2 “Washout” replicates (MB157, HCC1599, CUTLL1, IC8) it was selected as a real binding site. For ChIP-Seq analysis of histone modifications (H3K27ac, H3K4me1, H3K4me3, H3K18ac, H3K9ac) a peak had to be conserved in 3 out of 4 (DMSO/GSI or GSI/Washout) replicates. Peaks were filtered for ENCODE blacklisted regions downloaded from https://sites.google.com/site/anshulkundaje/projects/blacklists. Read counts per site were collected using the *summarzieOverlaps* function of the GenomicAlignments R/BioConductor package. DESeq2 was used to calculate normalization factors per replicate based on these read counts. DeepTools (33) *bamCoverage* function was used to calculate normalized BigWig files using the normalization factors provided by DESeq2 or the RPKM normalization for the remaining files. The BigWig files were used to visually inspect the quality of the ChIP-Seq experiments and the effectiveness of the peak calling in IGV (34).

DESeq2 was used for the identification of dynamic RBPJ binding sites and deregulation of ATAC-Seq and histone marks. In case of ATAC-Seq or histone modifications the window for detection of changes was from 500 bp upstream to 500 bp downstream of the RBPJ sites. All RBPJ binding sites with a log_2_FC < −0.5 upon GSI (or > 0.5 upon Washout of GSI) were selected as dynamic sites. DeepTools *computeMatrix* function was used to calculate a score per genome region matrix based on the normalized BigWig files and the binding sites as a reference. These matrices were used to plot the heat maps using deepTools *plotHeatmap* function or the average binding signal as line plots within R. For the identification of binding motifs at RBPJ sites the MEME-Suite v. 5.3.3 (35) was used. For the identification of head to head RBPJ binding motifs the *vmatchPattern* (36) function was used with the allowance of up to two mismatches. Position of the RBPJ binding sites in relation to the TSS was calculated and plotted using the ChIPseeker (37) package. Known mouse mm9 CpG islands were downloaded from UCSC table browser.

Annotation of RBPJ binding sites to their corresponding gene was performed using an in-house tool (that works in a comparable manner to basal plus extension of GREAT) in combination with the corresponding GTF file. Genes that are associated with both a dynamic and a static binding site were assigned as dynamic. The gene annotations were used for the gene over-representation analysis (GO & KEGG database) calculated with the clusterProfiler (38) package and plotted with ggplot2 (39).

### Prediction model

In order to predict dynamic and static RBPJ binding sites in Beko we used MSPC’s p-value (as a proxy for the quality of binding regions) and the position feature as defined by the ChIPseeker *annotatePeaks* function. Compensation for imbalance in the classifications was done by testing multiple different numbers (158, 500, 1000, 1500, 2500 and all) of static sites as input for the model. Therefore, random subsampling of static sites was used. The *createDataPartition* function of the caret (40) package was used to split the dataset (All dynamic sites + variable number of static sites) into 80% training data and 20% test data. The random forest model was generated based on the training data using the *randomForest* function of the randomForest (41) package. Mean accuracy (fraction of correctly predicted sites in test data) was measured over 500 random forest models based randomly sup-sampled static sites (Table S8). Finally, one random model with an accuracy over the mean of all 500 models was chosen as the final one. This model was tested and validated on the other cell types.

### Superenhancer

The BAM files of the two replicates for H3K27ac ChIP-Seq in Beko control cells were merged using samtools’ *merge* function. This merged BAM was used to call peaks using MACS2 (31) with the --nomodel --mm9”. The called peaks together with the merged BAM file was used as an input for the identification of superenhancer using ROSE (42).

### ATAC-Seq

ATAC-Seq was performed with the ATAC-Seq kit (Active Motif 53150) accordingly to manufactureŕs instructions and samples were sequenced on a NovaSeq device.

### ATAC-Seq analysis

Basic summary statistics for the ATAC-Seq experiments are provided in Table S6. ATAC-Seq primary analysis was performed identical to ChIP-Seq. MACS2 without input was used to for peak calling. Only peaks that were conserved in 3 out of 6 replicates were selected as real ATAC-Seq sites after removing blacklisted regions.

## RESULTS

### Identification of dynamic binding of transcription factor RBPJ

In order to study the genome-wide occupancy of TF RBPJ, we performed ChIP-Seq analysis in a mouse progenitor T-cell line called Beko (6,16). Beko cells are constitutively active for Notch signaling and formation of NICD1 can be blocked by treatment with γ-secretase inhibitor (GSI; Figure 1A). In control DMSO-treated Beko cells, a total of 3558 RBPJ binding sites were detected (Table S1). This number of binding sites is comparable to previously published data in mouse mature T-cells (43). When we compared the RBPJ binding profile in presence or absence of GSI we observed that the majority of sites (3380) remained unaffected by GSI treatment, but a small fraction (158 sites) showed a marked reduction of RBPJ binding (log_2_ fold change cut-off < −0.5) upon addition of GSI; we refer to these sites as either static or dynamic sites, respectively (Figure 1B-C and Table S1). Interestingly, the RBPJ binding is stronger at dynamic sites compared to static sites when the Notch pathway is active (Figure 1C). Differences in binding strength could be validated by ChIP-qPCR (Figure S1A) and also canonical RBPJ motif ‘TGGGAA’ were observed (Figure S1B and Table S2). The RBPJ motif identified at dynamic sites had a less conserved ‘G’, but both purines occurred with equal frequency (TGRGAA). Both static and dynamic sites are enriched for binding motifs for members of the SP (Specificity Protein) family (Figure S1B and Table S2). Interestingly, the SP motif is predominantly identified at static sites, whereas the RBPJ motif marks a large fraction of the dynamic sites (Figure S1B and Table S2). In addition, static sites are enriched for NRF1, NFYA/NFYB/DUX, FOXJ3/ZF384 and ZBTB7A/GABPA/ELK4 motifs while dynamic sites are enriched for the TCF3/TCF4 motif (Table S2). Previous studies have described that Notch target genes are regulated by dimeric RBPJ complexes that bind to dimeric DNA binding motifs oriented head-to-head and with a distance of 15-17 nucleotides (1,44–47). Interestingly, we observed that dimeric RBPJ binding sites are preferentially enriched within the group of dynamic sites (Figure S1C).

**Figure 1.**
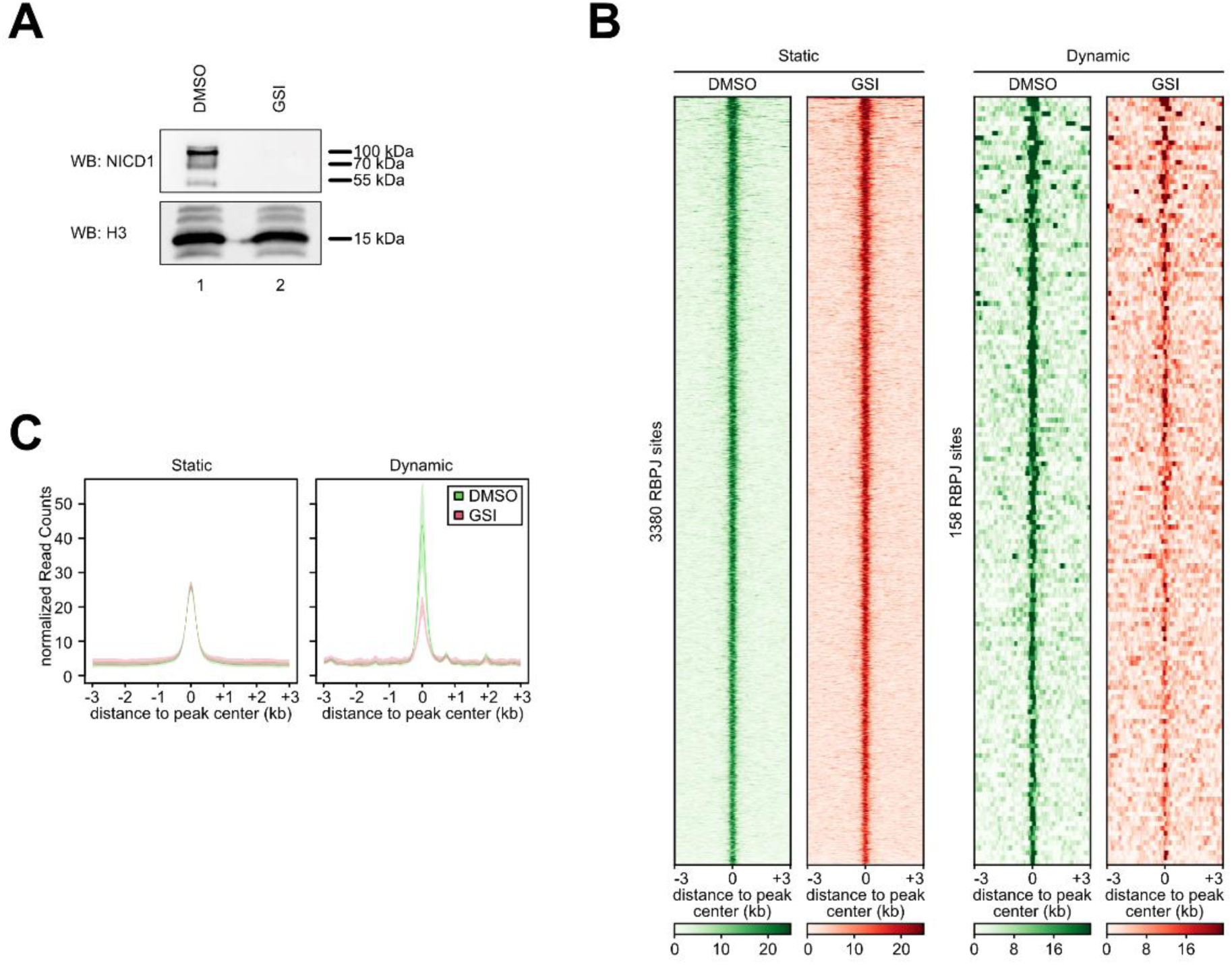
Identification of static and dynamic RBPJ binding sites in Beko cells. (**A**) The active cleaved NICD1 protein disappears after GSI treatment for 24 hours. Beko cells were treated for 24 hours with 10 μg/ml GSI or DMSO as control and the nuclear extracts (NE) were analyzed by Western blotting versus the endogenous cleaved Notch1 protein (NICD1) or H3 as loading control. (**B**) Heat map showing the static and dynamic RBPJ binding sites identified in control Beko cells and their behavior upon treatment with GSI. Beko cells were treated for 24 hours with 10 μg/ml GSI or DMSO as control and the genome-wide binding of RBPJ was investigated by ChIP-Seq. (**C**) Line plots showing the average RBPJ binding signal for static and dynamic sites. Outline depicting the standard deviation of the replicates Beko cells were treated for 24 hours with 10 μg/ml GSI or DMSO as control and the genome-wide binding of RBPJ was investigated by ChIP-Seq.

Taken together, our results in Beko cells show that RBPJ occupancy is strongly reduced at only a small fraction of binding sites upon pharmacological inhibition of the Notch pathway.

### Characterization of dynamic RBPJ binding sites

Next, we characterized the differences between static and dynamic RBPJ binding sites, taking genomic and chromatin features into account. Most static sites are located close to transcriptional start sites (TSS; Figure 2A and S2A). In contrast, dynamic sites were preferentially found in intra- and intergenic regions (Figure 2A and S2A). Both static and dynamic sites were associated with open chromatin regions as measured by ATAC-Seq (Figure 2B). Overall, the accessibility was higher at static RBPJ sites compared to dynamic ones (Figure 2B). In addition, both groups of RBPJ sites were enriched for active chromatin mark H3K27ac (Figure 2C). In line with the differences in binding position (Figure 2A), enhancer mark H3K4me1 is highly enriched at dynamic sites whereas H3K4me3, predominantly found at TSSs, was highly enriched at static sites (Figure 2D-E, respectively). In addition, RBPJ binding highly overlaps with CpG islands (CGI; Figure S2B) and closer inspection unveiled that static RBPJ sites are more associated with CGI than dynamic sites (Figure S2C), in line with the notion that promoters are CGI-rich (48,49). Of note, the higher enrichment of H3K4me3 at static sites further reflects their co-localization at promoters, which has been previously described (50). We have previously described the role of the lysine methyltransferase 2D (KMT2D) in the dynamic regulation of H3K4me3 in Notch target gene activation (9). Therefore, we focused on this mark to have a better genome-wide view of its regulation in response to the Notch signaling. We observed that static H3K4me3 sites that overlap with RBPJ sites are closer to the TSS compared to dynamic H3K4me3 sites that overlap with RBPJ (Figure S2D). In addition, co-localization of H3K4me3 sites and RBPJ sites overlap more frequently with CGI compared to dynamic H3K4me3 sites overlapping with RBPJ (Figure S2E). These data suggest that H3K4me3 is more dynamic at distal than promoter proximal RBPJ binding sites, in line with our previous observations (9).

**Figure 2.**
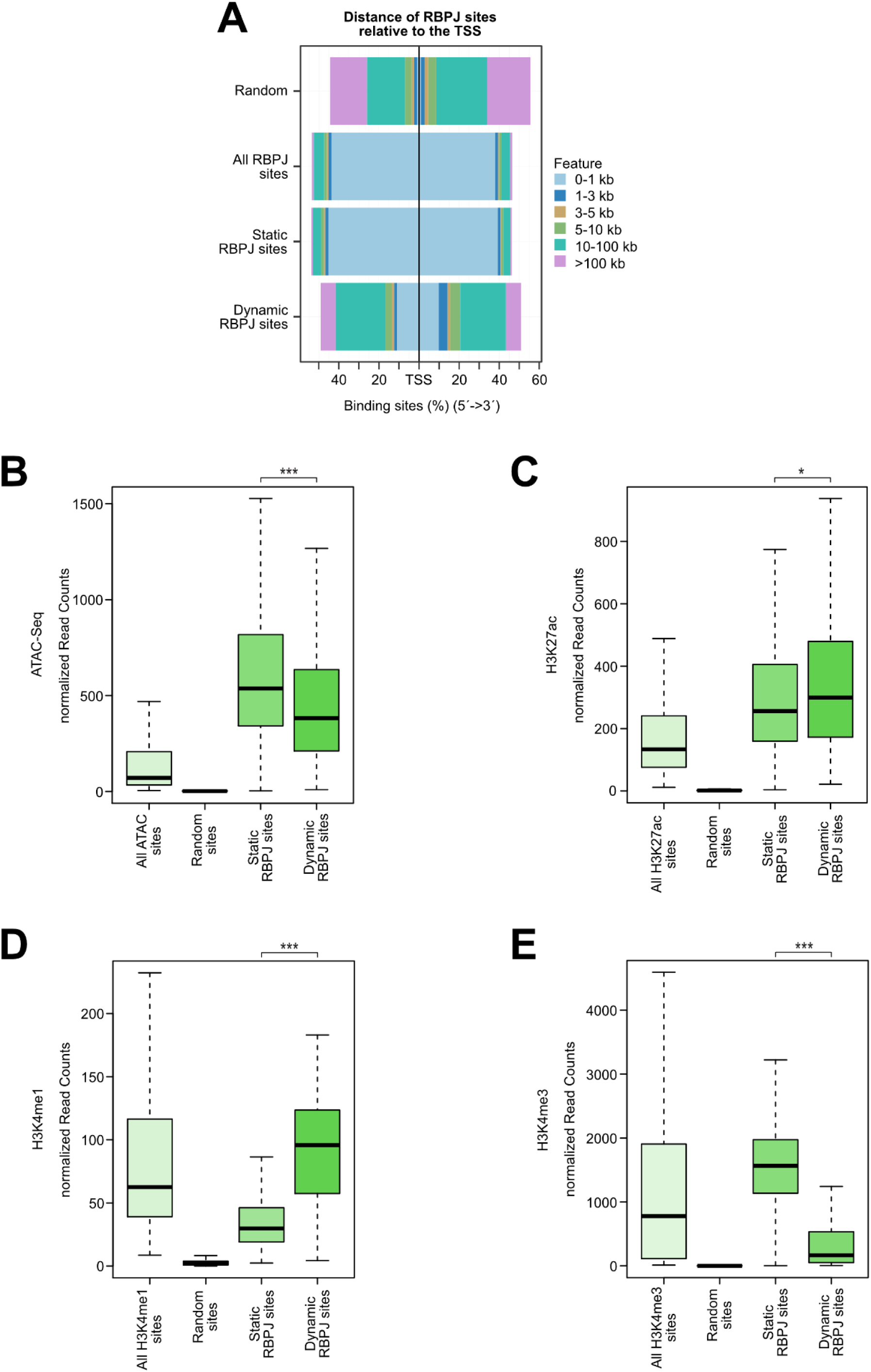
Dynamic RBPJ sites represent distal enhancers. (**A**) Distance of all, static and dynamic RBPJ sites to the next transcription starting site (TSS). The figure includes the genomic background distribution (random). (**B-E**) Beko cells were analyzed by ATAC-Seq and ChIP-Seq to characterize the chromatin landscape at static and dynamic RBPJ sites or at random sites and all detected sites of the given mark (or ATAC-seq) as control. (**B**) Box plot showing that both dynamic and static sites display an open chromatin configuration as measured by ATAC-Seq. The chromatin accessibility is higher at static sites compared to dynamic ones. (**C**) Box plot showing that H3K27ac is higher at dynamic compared to static RBPJ sites. (**D**) Box plot showing that H3K4me1, a typical enhancer mark, is higher at dynamic compared to static RBPJ sites. (**E**) Box plot showing that H3K4me3 is higher at static compared to dynamic RBPJ sites. Wilcoxon rank sum tests (\*\*\**P <* 0.001, \**P <* 0.05).

In conclusion, dynamic and static RBPJ binding sites reveal both high level of chromatin accessibility and activity but differ in regard to distance to the TSS, which is reflected in chromatin marks.

### Transcriptional response to Notch activation is preferentially associated with dynamic sites

In order to investigate the functional consequences of dynamic RBPJ binding we associated the RBPJ sites to nearby genes and we investigated whether those genes were influenced by GSI treatment in Beko cells. Genes associated with dynamic RBPJ sites were significantly downregulated, while genes associated with static sites were not affected (Figure 3A and Table S3). In support of that, we observed that dynamic sites are statistically more frequently associated with deregulated genes than expected by chance (Figure S3A).

**Figure 3.**
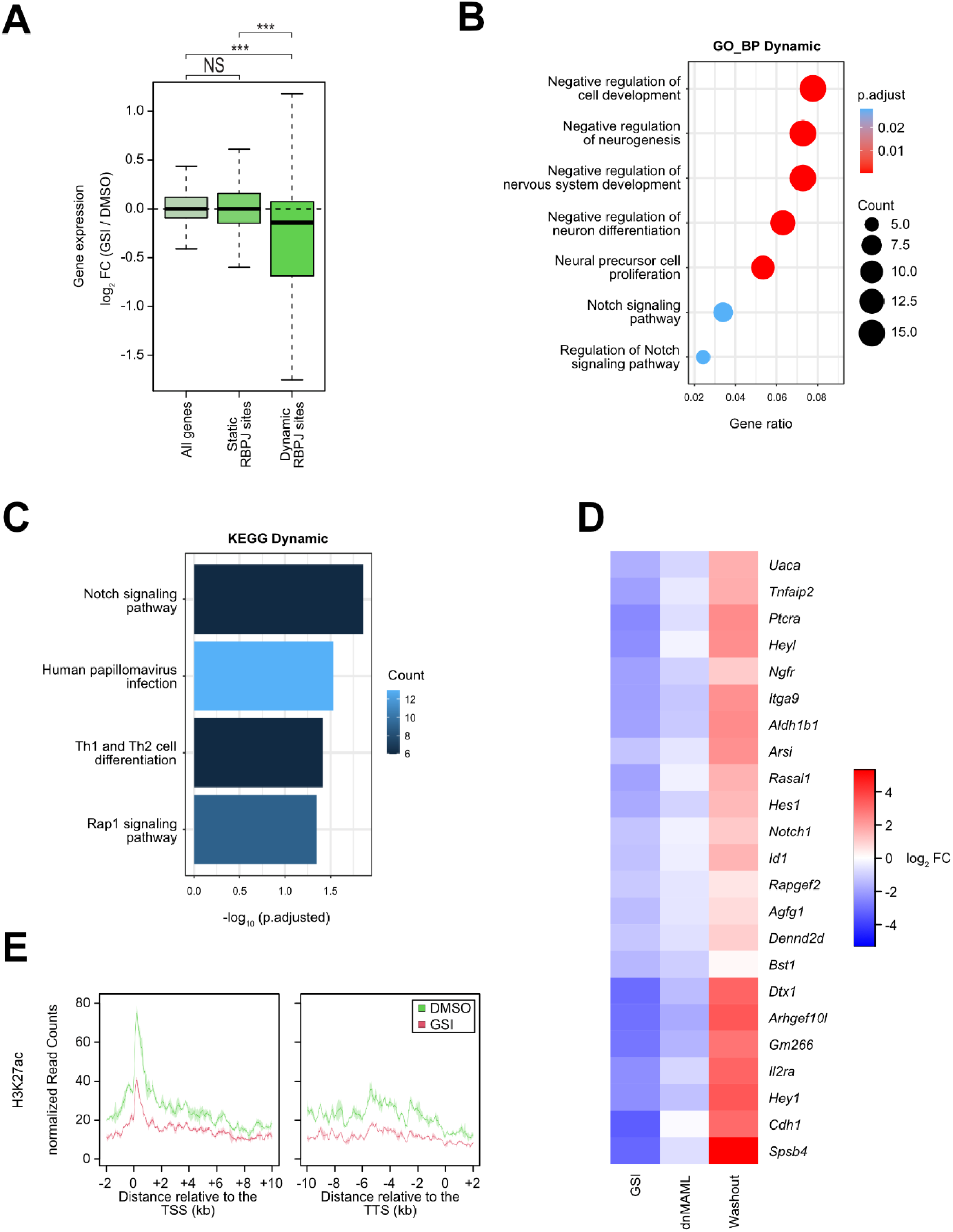
Transcriptional Notch responsiveness is associated with dynamic RBPJ sites. Beko cells were treated for 24 hours with 10 μg/ml GSI or DMSO as control. (**A**) Box plot showing the effects of GSI treatment on the expression of genes associated with static or dynamic RBPJ sites. Wilcoxon rank sum tests (\*\*\**P <* 0.001, NS = not significant). (**B**) ORA based on the gene ontology (GO) database for the group of dynamic RBPJ sites identified in Beko cells. The panel depicts the results for the “Biological Process” (BP) class. Full list of the GO terms identified in this study is available in Table S4. (**C**) ORA analysis based on the KEGG database for the group of dynamic RBPJ sites identified in Beko cells. Full list of the KEGG terms identified in this study is available in Table S4. (**D**) Heat maps showing the effects of a dominant negative mutant of MAML (dnMAML) (60) or of GSI washout on the expression of genes downregulated by GSI and associated with dynamic RBPJ binding sites. (**E**) Line plot showing the effects of GSI treatment within the gene bodies of those genes significantly downregulated upon GSI and associated with dynamic RBPJ sites on H3K27ac as measured by ChIP-Seq. Outline depciting the standard deviation of the replicates TSS: Transcription starting site; TTS: Transcription termination site.

Subsequently, we examined the functions of genes associated with either static or dynamic RBPJ sites. To this end, we analyzed these genes using an overrepresentation analysis (ORA) to test if they were enriched for genes associated with different known biological pathways. The genes associated with dynamic sites were enriched for different Notch-associated terms from gene ontology (GO) and KEGG databases (Figure 3B-C and Table S4). Furthermore, GO terms associated with negative regulation of development and neurogenesis pathways were significantly enriched within the genes with dynamic RBPJ sites (Table S4). The term “Th1 and Th2 cell differentiation” was significantly enriched, when using the KEGG database. This is in line with previous publications, which highlighted the important role of Notch signaling in T helper cell development (51). In contrast, the genes associated with static sites were only enriched for one Notch associated term (“Notch receptor processing”) based on the GO and KEGG databases (Figure S3B-C). We also observed that significantly downregulated genes associated with dynamic RBPJ sites are also downregulated upon overexpression of a dominant negative mutant of the Notch cofactor Mastermind-like (dnMAML) (Figure 3D). We further validated our findings by performing a washout of GSI: We treated Beko cells for 48 hours with GSI and after that, we washed out (washout) the inhibitor and placed the cells back in culture for additional 24 hours before analyzing their gene expression profile. First of all, we observed that the cleaved active NICD1 protein reappears in the nuclear fraction upon GSI washout (Figure S4A). The washout experiment matches quite well with our previous GSI gene expression data (16). In fact, genes downregulated by GSI were upregulated upon washout and genes upregulated by GSI are downregulated upon washout (Figure S4B-E and Table S4). Importantly, genes associated with dynamic RBPJ sites are preferentially upregulated upon GSI washout (Figure S4F). Similarly, genes downregulated by GSI and associated with dynamic RBPJ sites are strongly upregulated upon GSI washout (Figure 3D).

Having identified genes downregulated upon GSI and associated with dynamic RBPJ sites, we next investigated the effects on the chromatin configuration of those genes. We observed reduced H3K27ac and H3K4me3 upon GSI treatment (Figure 3E and S5). However, chromatin accessibility was hardly affected at both the TSS and within the gene bodies of genes downregulated by GSI and associated with dynamic RBPJ site(s) Figure S5).

Altogether, these data suggest that genes associated with dynamic RBPJ sites are preferentially regulated by the Notch signaling pathway and that the effects of this regulation are reflected on the chromatin dynamics.

### Dynamic RBPJ sites are associated with superenhancers

In order to further characterize the Notch transcriptional complexes, we focused on the analysis of chromatin configuration in response to inactivation of Notch signaling at RBPJ sites. We observed reduced chromatin accessibility, H3K27ac and H3K4me3 while H3K4me1 was unaffected at dynamic RBPJ sites in response to GSI treatment (Figure 4A-D). On the other side, minor but statically significant changes were observed at static RBPJ sites in response to inhibition of Notch signaling (Figure 4A-D). Previous works highlighted a dynamic regulation of superenhancers (SEs) in response to Notch signaling (Wang et al. 2014). In conjunction with the H3K27ac ChIP-Seq data, we used the ROSE tool and detected 935 SEs, which is a comparable number to previously identified SE in other cells (52). As expected, H3K27ac is strongly enriched over the entire span of the identified SEs (Figure S6). In addition, we observed that H3K4me1- and to a lesser extend H3K4me3 marks are enriched over the entire length of the identified SEs (Figure S6). Having identified the SEs in Beko cells, we proceed to characterize them in relation to Notch responsiveness. We first observed that a much larger fraction of dynamic RBPJ sites overlap with SEs, compared to static RBPJ sites (Figure 4E). Furthermore, we observed that H3K27ac is preferentially reduced at SEs that contain dynamic RBPJ sites compared to the ones that have static RBPJ sites in response to perturbation of Notch signaling (Figure 4F). Together, these data suggest that Notch responsiveness is associated with dynamically regulated RBPJ sites and these dynamic RBPJ sites often localize within SEs.

**Figure 4.**
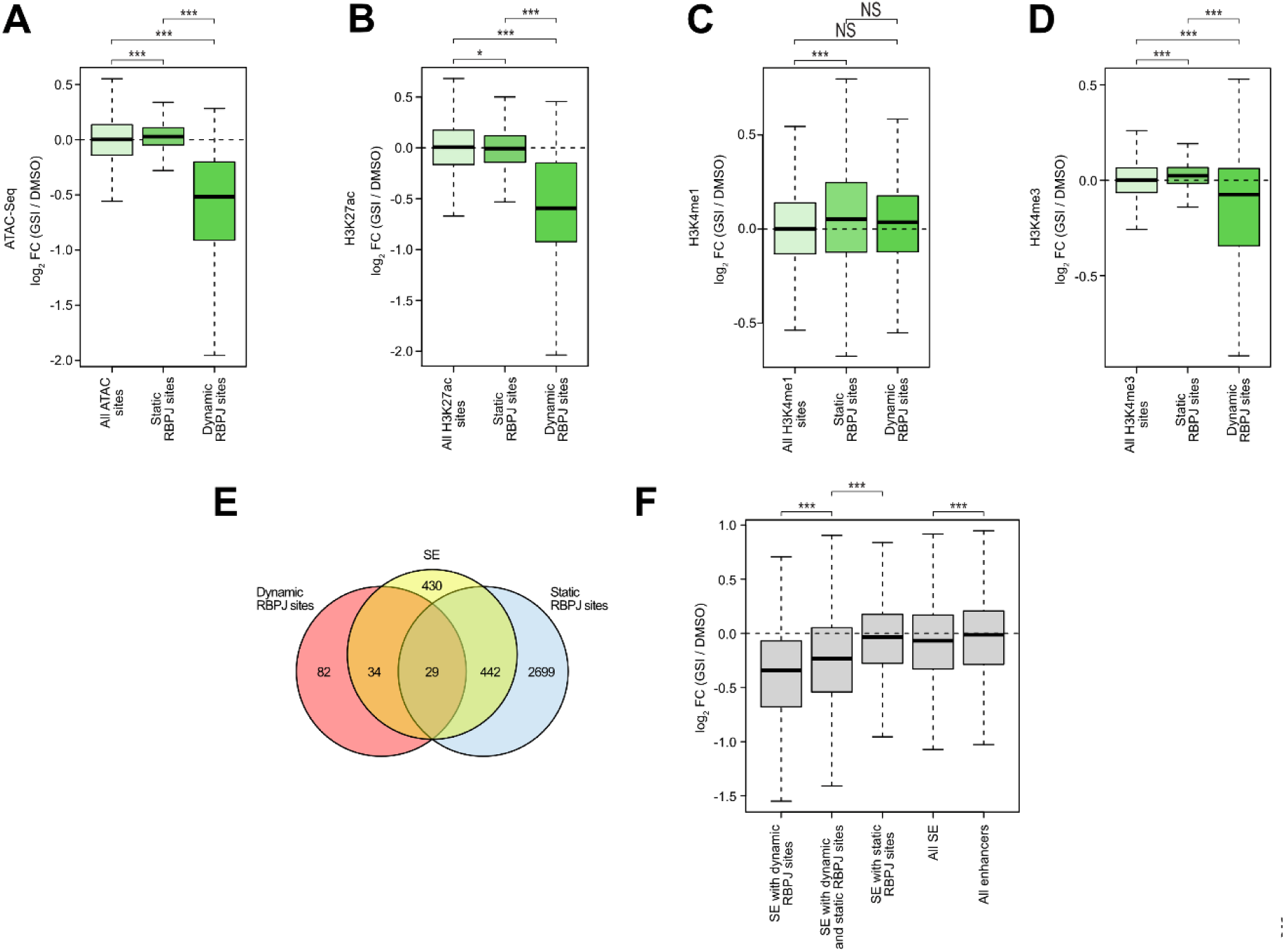
Notch responsiveness on the chromatin is associated with dynamic RBPJ sites. Beko cells were treated for 24 hours with 10 μg/ml GSI or DMSO as control. Changes on the chromatin configuration were analyzed by ATAC-Seq and ChIP-Seq (**A-D**). Box plot showing the effects of GSI treatment on (**A**) chromatin accessibility, (**B**) H3K27ac, (**C**) H3K4me1 and (**D**) H3K4me3 at static and dynamic RBPJ sites as measured by ATAC-Seq and ChIP-Seq. Wilcoxon rank sum tests (\**P <* 0.05, \*\*\**P <* 0.001, NS = not significant). (**E**) Venn diagram showing the overlap of superenhancers (SEs) with dynamic or static RBPJ sites in Beko cells. (**F**) Boxplot showing the changes in H3K27ac sites within all superenhancers, SEs associated with dynamic RBPJ sites, with static RBPJ sites or with both dynamic and static RBPJ sites.

### Dynamic binding behavior is conserved in triple-negative breast cancer cells

As a next step, we investigated whether the dynamic RBPJ binding is a predictive characteristic of Notch responsiveness in other cell types. To reach this goal we analyzed publicly available datasets from triple negative breast cancer (TNBC) cells (HCC1599 and MB157), in which the Notch signaling pathway has been first blocked with GSI and subsequently reactivated by washout of the inhibitor (19). TNBC is a subtype of breast cancer that is characterized by the lack of ER, PR, HER2 and is known to have active Notch signaling (53). Using RBPJ ChIP-Seq, we identified 14010 binding sites in HC1599 and 7628 in MB157, respectively (Figure S7A and S8A, respectively). In line with our findings in Beko cells, in both HCC1599 and MB157 cells static and dynamic binding behavior of RBPJ upon washout of GSI was detectable (Figure S7A and S8A, respectively). In HCC1599, 2607 (~18.6%) of the 14010 RBPJ sites had increased RBPJ binding after washout of GSI (Figure S7A). In MB157, 2040 of 7628 (~26.7%) sites were dynamically bound in response to reactivation of Notch signaling (Figure S8A). Having identified dynamic and static RBPJ binding sites in HCC1599 and MB157 cells, we proceeded with their characterization.

In Beko cells, static sites are preferentially enriched at promoter proximal regions, whereas dynamic sites are mostly located at distal regulatory elements (Figure 2A and S2A). In order to test whether this binding behavior is characteristic also of TNBCs, we tested the binding position of dynamic and static RBPJ sites relative to the next TSS in both HCC1599 and MB157. Static RBPJ sites localize much closer to the TSS than the dynamic ones in both cell lines (Figure S7B and S8B). Subsequently, we evaluated whether Notch responsiveness on the chromatin level, measured via H3K27ac levels, is preferentially associated with dynamic RBPJ sites. We observed significant changes in H3K27ac preferentially at dynamic RBPJ sites compared to the static ones upon perturbation of Notch signaling (Figure S7C and S8C).

Our results in Beko cells suggest that binding strength and positioning of the RBPJ sites are good predictors of Notch responsiveness. Based on these results, we ranked the RBPJ sites in HCC1599 and MB157 by the quality of the peaks as measured by their respective p-values. We observed that a high ranking position is strongly correlated with dynamic binding of RBPJ (Figure 5A and D). We observed the inverse trend when looking at the distance to the TSS, where the fraction of dynamic sites becomes smaller with decreasing distance to the TSS (Figure 5B and E). In order to investigate whether there is a connection between Notch-dependent transcriptional response and dynamic sites also in TNBC cells, we tested the statistical enrichment of deregulated genes within the groups of genes that are associated with static or dynamic RBPJ sites. Very similar to Beko cells, we observed a higher enrichment of deregulated genes within the group of dynamic sites (Figure 5C and F).

**Figure 5.**
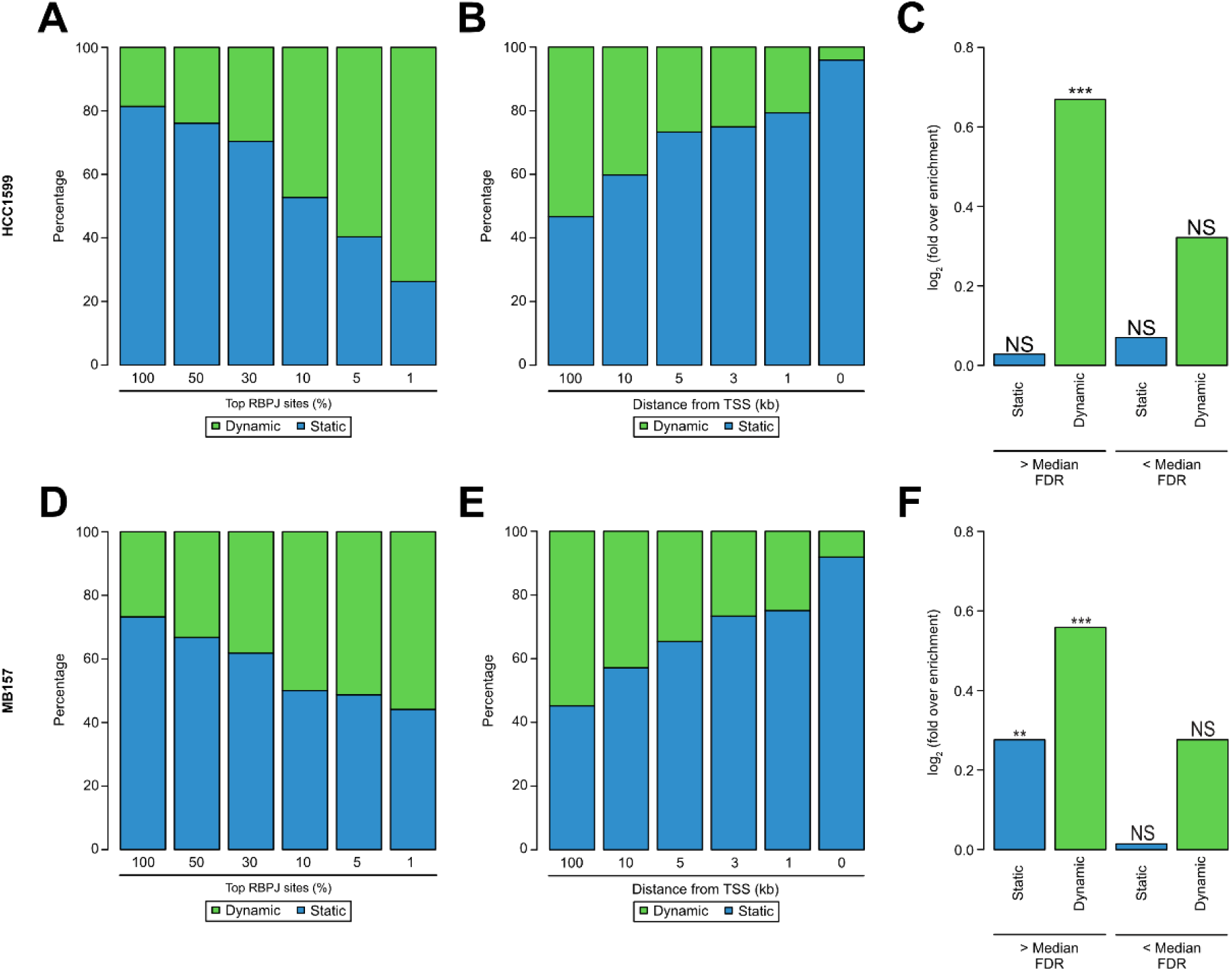
Notch responsiveness is preferentially associated with dynamic RBPJ sites in triple negative breast cancer (TNBC). Publicly available ChIP-Seq and RNA-Seq data were analyzed to investigate Notch responsiveness in HCC1599 (**A-C**) and MB157 (**D-F**) triple negative breast cancer (TNBC) cells. (**A and D**) Bar plot showing the correlation between static and dynamic RBPJ binding sites and their respective peak quality (MSPC’s p-value) in HCC1599 cells (**A**) or MB157 cells (**D**). (**B and E**) Bar plot showing the correlation between static and dynamic RBPJ binding sites and their distance to the transcription starting site (TSS) in HCC1599 cells (**B**) or MB157 cells (**E**). (**C and F**) Bar plot showing the enrichment of significantly deregulated genes within genes associated with only static or only dynamic RBPJ sites in HCC1599 cells (**C**) or MB157 cells (**F**). Additionally, the top half (> Median) and the bottom half (< Median) of all RBPJ sites shown. In these cases, we assigned genes associated with both static and dynamic sites as dynamic. Hypergeometric test (\*\**P <* 0.01, \*\*\**P <* 0.001, NS = not significant).

Taken together, these results suggest that dynamic RBPJ sites are characterized by being mostly located far away from TSS as well as being enriched for the high-quality sites in Beko cells and in TNBCs.

### Predicting dynamic and static RBPJ sites

Finally, we used a machine learning algorithm to predict dynamic and static RBPJ sites. This is potentially useful to further refine and validate the conserved characteristics of dynamic and static binding sites taking a potentially unbiased approach. Moreover, this could also allow us to identify cell type-specific Notch target genes in any given cell-type.

As our training set, we used data from Beko cells since they represent our best described system. We took all 158 dynamic sites and only used 1500 random static sites to generate the training set used for the prediction model (Figure 6A). A random forest using the normalized p-value calculated of the peak and the genomic feature as a training set led to the best prediction of static and dynamic RBPJ sites in Beko cells (Figure 6A-B and G). The model could efficiently predict both static and dynamic RBPJ sites (Figure 6B, F, G and Table S6). In order to test whether the model created is actually able to predict dynamic and static sites in other cell types, we first tested the model on the two TNBC cell lines described above. We were able to predict most of the static and dynamic sites in HCC1599 (Figure 6B-G and Table S6) and in MB157 (Figure 6D, G and Table S6).

**Figure 6.**
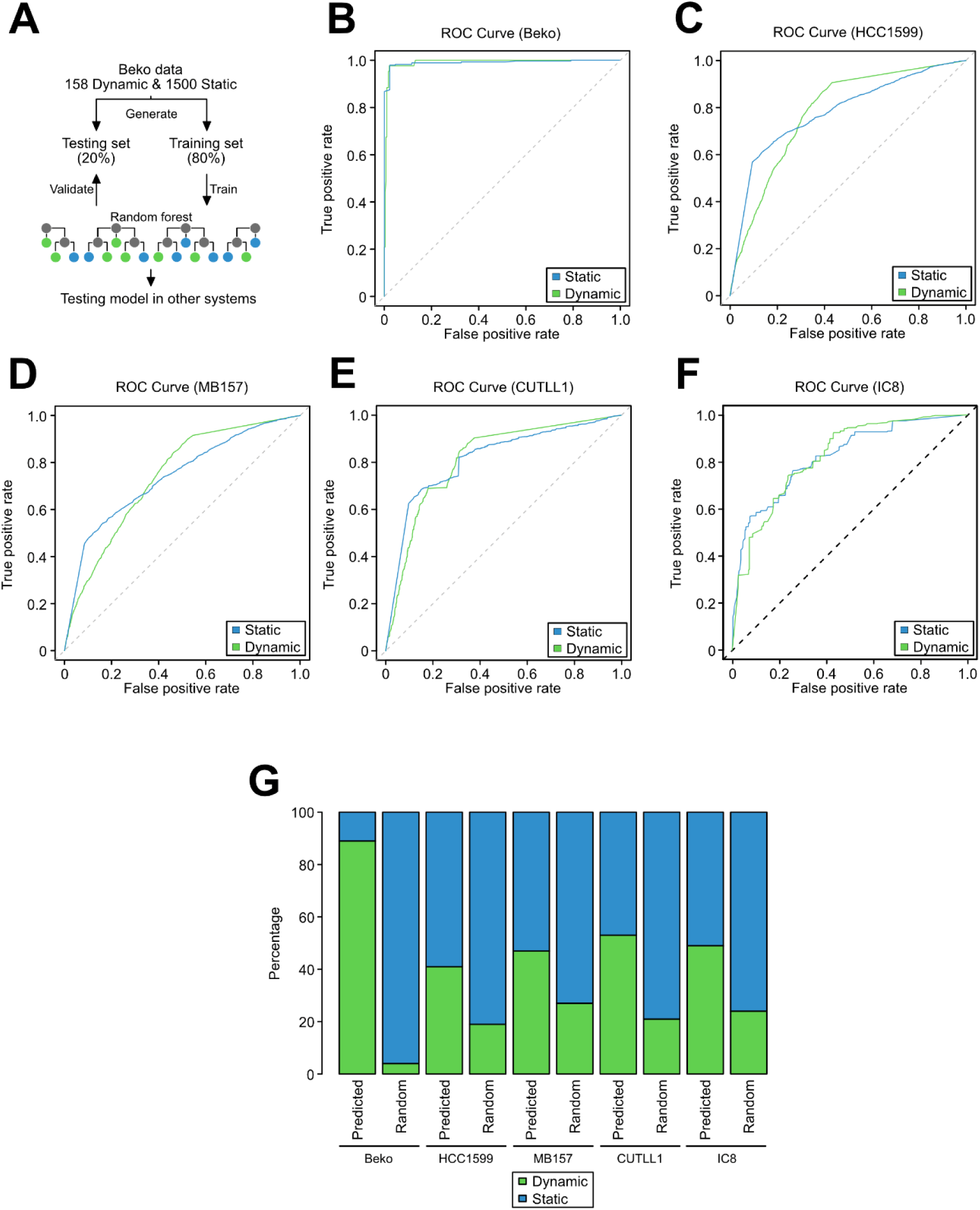
Development of a model for prediction of Notch responsiveness. (**A**) Schematic representation of the approach used to develop and test the prediction model used to identify static and dynamic RBPJ binding sites. All the 158 dynamic RBPJ sites and a random selection of 1500 static RBPJ sites were used to generate a random forest model. We used 80% of the selected RBPJ sites (both static and dynamic) to train the model whereas the residual 20% were used for testing. (**B**) ROC curves showing the true and false positive rates of the prediction model for static and dynamic sites independently in Beko cells. (**C-F**) The prediction model is able to identify static and dynamic RBPJ sites with high accuracy in HCC1599, MB157, CUTLL1 and IC8 cells. (**G**) Bar plot comparing predicted versus randomly selected identifying dynamically bound RBPJ binding sites in Beko, HCC1599, MB157, CUTLL1 and IC8 cells using the machine learning approach.

Finally, we searched for other testable data sets and used data from two other cell lines: Human T cell acute lymphoblastic leukemia (T-ALL) called CUTTL1 and human squamous cell carcinoma (SCC) called IC8, both also employing washout of Notch inhibitor GSI to dynamically regulate the Notch response (21,24,54). We could efficiently identify responsive RBPJ sites in CUTTL1 (Figure 6E, G and Table S6) and in IC8 (Figure 6F, and Table S6). In summary, our machine learning approach allowed us to efficiently and consistently identify more RBPJ dynamic sites than expected by chance (Figure 6G).

To further validate the efficiency of the predicted dynamic and static sites, we compared the actual measured changes in RBPJ binding upon GSI washout for the predicted static or dynamic RBPJ sites. Worth mentioning, the criterion to define an RBPJ site as dynamic depended to changes in Notch activity (log_2_FC > 0.5 for the washout of GSI or < −0.5 for the GSI treatment). In HCC1599 (Figure 7A), MB157 (Figure 7B) and CUTLL1 (Figure 7C) cell lines significant differences between the predicted static and dynamic sites were also detectable. For these cell lines, the predicted dynamic sites had a much stronger increase of RBPJ binding upon washout of GSI compared to the predicted static sites.

**Figure 7.**
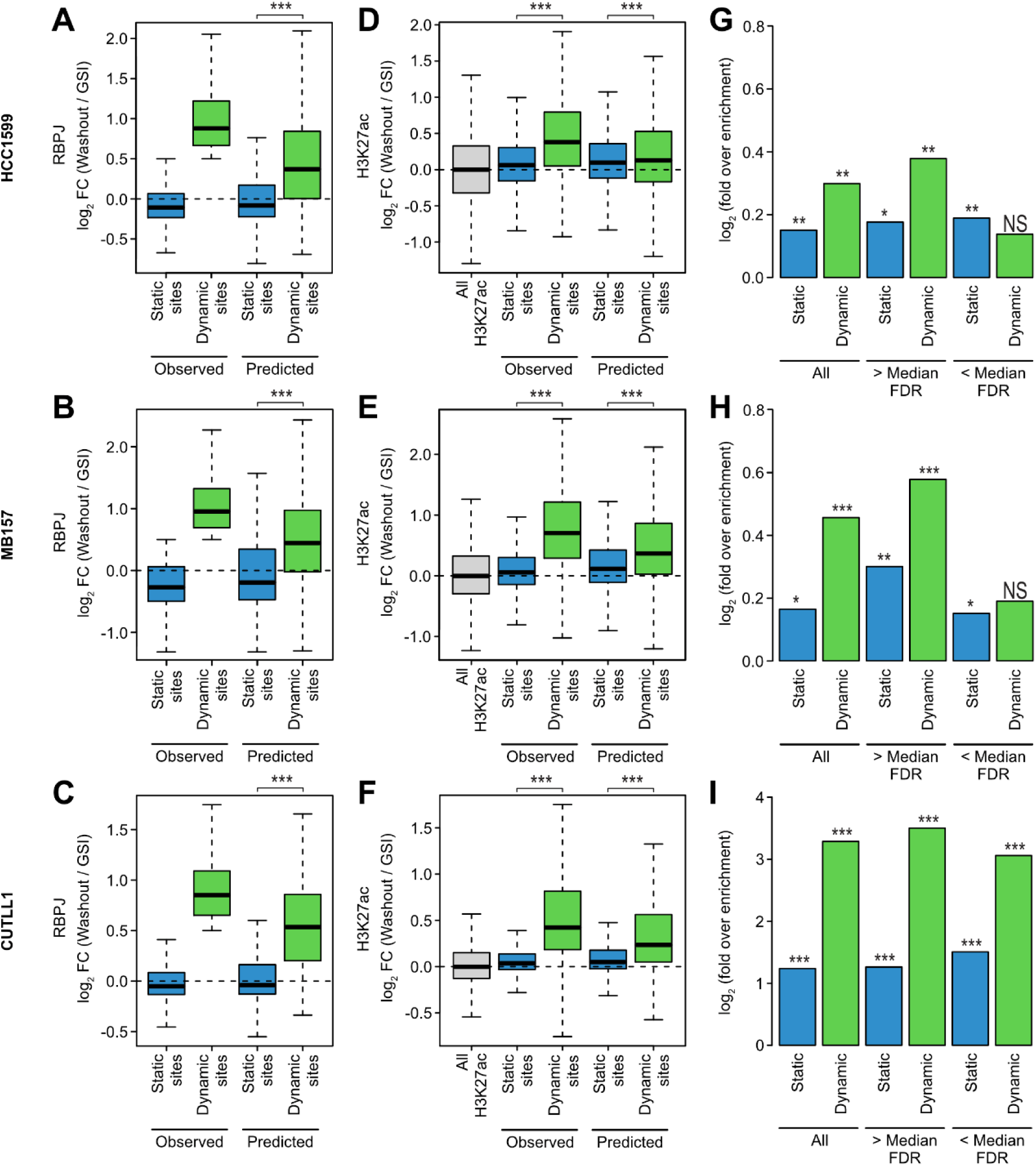
Predicted dynamic sites are comparable to observed ones. (**A-C**) Box plots showing the RBPJ binding changes upon GSI washout for observed and predicted static or dynamic RBPJ binding sites in HCC1599 (**A**), MB157 (**B**) and CUTLL1 (**C**) cells. (**D-F**) Box plots showing the H3K27ac changes upon GSI washout for observed and predicted static or dynamic RBPJ binding sites in HCC1599 (**D**), MB157 (**E**) and CUTLL1 (**F**) cells. (**G-I**) Bar plot showing the enrichment of significantly deregulated genes within genes associated with predicted static or dynamic RBPJ sites in HCC1599 (**G**), MB157 (**H**) and CUTLL1 (**I**) cells. Additionally, the top half (> Median) and the bottom half (< Median) of all RBPJ sites are shown. Here are genes associated with only static or dynamic (and static) RBPJ sites. Hypergeometric test (\**P <* 0.05, \*\**P <* 0.01, \*\*\**P <* 0.001, NS = not significant).

An additional feature of dynamic sites was the changes in H3K27ac levels. In order to test, whether these changes were still detectable for the predicted static and dynamic sites, we analyzed H3K27ac ChIP-Seq upon washout of GSI in HCC1599, MB157 and CUTLL1. As expected, there is a significant increase in H3K27ac levels at the dynamic sites compared to the static ones in both HC1599 (Figure 7D), MB157 (Figure 7E) and CUTLL1 (Figure 7F). Subsequently, we analyzed the transcriptional response of genes associated with predicted static and dynamic RBPJ sites. The previous analysis showed that genes associated with dynamic sites were more likely significantly differentially regulated in response to Notch signaling. To this end, we used the beforementioned enrichment of differentially expressed genes (DEG) analysis for the groups of predicted static and dynamic RBPJ binding sites. In line with our results in Beko cells, the enrichment of DEG was much higher for predicted dynamic sites than for predicted static sites for all cell lines under investigation (Figure 7G-I). Strikingly, for CUTLL1 the weaker predicted RBPJ sites [< median false discovery rate (FDR) of the corresponding RBPJ peak] had a stronger enrichment for DEG compared to the strongest predicted static sites (> median FDR of the RBPJ peak). Therefore, strong binding, despite being strongly correlated with dynamic sites, is not sufficient to classify RBPJ binding site behavior. This indicates that the predictive power of the model is higher than its individual input features.

## DISCUSSION

Here, we propose a model for the responsiveness of Notch target genes solely based on the localization and binding strength of the TF RBPJ. We are able to predict with high confidence functional target sites and thus Notch responsiveness.

When focusing on the differences of sequence context between dynamic and static sites, it is surprising that the dynamic sites carry the canonical RBPJ binding motif ‘TGGGAA’ far more often than the static ones. This could indicate that at dynamic sites RBPJ binds directly to DNA, whereas at static sites the binding may have to be stabilized by additional factors. At the same time these factors may mask certain accessible cofactors or chromatin features and thus prevent activation. We have seen that RBPJ motifs at dynamic and static sites are characterized by subtle differences. Moreover, static RBPJ sites are also marked by additional binding motifs and it remains to be seen whether these contribute to the lack of responsiveness, which may point to a commonly used mechanism. Interestingly, one of top motifs identified at static sites is the Ronin/HCF-1 motif (55,56). This has been described to be specifically associated with housekeeping promoters that are sufficient for robust expression of genes lacking distal enhancers. We speculate that these promoters are particularly robust to perturbations, Noteworthy, the Ronin/HCF-1 motif is composed of two small motifs separated by 2-3 nucleotides, one of which being extremely similar to the RBPJ motif. This might indicate either imply that TFs compete or form a Notch-non-responsive configuration.

In line with this, the vast majority of static sites is associated with transcriptional start sites. It has been described that many promoters are resistant to transcriptional perturbations and are tuned to transcriptional robustness. Interestingly, even the shape of the promoter and number of TF binding sites dictate the plasticity of promoters (57,58). Thus, RBPJ binding even together with a coactivating complex might not be decisive in the context of such broad promoters with multiple competing TFs. In contrast to the static sites, a characteristic of a dynamic RBPJ sites could be that the chromatin accessible region is narrow and can possibly form chromatin loops. Apart from looping presence of CTCF sites might also influence the potentially dynamic binding of RBPJ. (19,59).

Taken together, it is conceivable that dynamic RBPJ sites represent the well-known Notch targets, while RBPJ at static sites could reflect spurious or indirect binding, which could for example serve as a ‘gate-keeper’ function.

The heterogeneity of the transcriptional outcomes of Notch activation remains a major conundrum. One proposition is that the activation complex consisting of RBPJ / NICD and additional coactivators has enhanced binding activity or that two such coactivator complexes bind cooperatively and this requires two head-to-head RBPJ sites (46,47). Such sites do not account for the majority of dynamic sites determining Notch responsiveness. Our data indicates that enhancer positioning relative to the TSS is more decisive. Our machine learning approach, focusing on RBPJ binding sites and genomic features, not only reduces the time and effort to identify robust Notch responsive genes with good accuracy, but also suggests that this could apply for other TFs factors, such as p53 or TCF of the Wnt signaling pathway. Our data strongly supports the notion that for RBPJ and Notch target genes a locus control region or superenhancer, is important for gene responsiveness. In future, it will be interesting to investigate whether this is also the case for other inducible TFs.

Taken together, computational analyses of TF RBPJ binding combined with distinct genomic features can be used to identify Notch responsiveness in any given cell type. Likely, a comparable model can be established for other inducible systems.

## COMPETING INTEREST STATEMENT

The authors declare no conflict of interest.

## ACKNOWLEDGEMENTS

We are grateful to T. Schmidt-Wöll for excellent technical assistance. The authors wish to acknowledge Centro de Análisis Genómico (CNAG-CRG), Spain, for sequencing the ChIP samples.

This work was funded by a research grant of the University Medical Center Giessen and Marburg (UKGM) and by a Prize of the Justus Liebig University Giessen to B.D.G. T.B. is supported by the Deutsche Forschungsgemeinschaft (DFG, German Research Foundation) - TRR81-A12 and BO 1639/9-1, the Behring-Röntgen foundation and Excellence Cluster for Cardio Pulmonary System (ECCPS) in Giessen. Funding for open access charge was provided by the DFG collaborative research TRR81.

## Author contributions

F.F. and B.D.G. performed experiments. T.F. performed the bioinformatics analysis. B.D.G., T.F. and T.B. conceived the study. B.D.G., M.B. and T.B. supervised the work. B.D.G., T.F. and T.B. wrote the manuscript with contributions from other authors.

## Notes

### Competing Interest Statement

The authors have declared no competing interest.

